# Voltage-dependent reversal potentials in spiking recurrent neural networks enhance energy efficiency and task performance

**DOI:** 10.1101/2025.08.29.673001

**Authors:** Miguel Rodrigues, Carmen Gasco-Galvez, Martin Vinck

## Abstract

Spiking recurrent neural networks (SRNNs) rival gated RNNs on various tasks, yet they still lack several hallmarks of biological neural networks. We introduce a biologically grounded SRNN that implements Dale’s law with voltage-dependent AMPA and GABA reversal potentials. These reversal potentials modulate synaptic gain as a function of the postsynaptic membrane potential, and we derive theoretically how they make each neuron’s effective dynamics and subthreshold resonance input-dependent. We trained SRNNs on the Spiking Heidelberg Digits dataset, and show that SRNN with reversal potentials cuts spike energy by up to 4×, while increasing task accuracy. This leads to high-performing Dalean SRNNs, substantially improving on Dalean networks without reversal potentials. Thus, Dale’s law with reversal potentials, a core feature of biological neural networks, can render SRNNs more accurate and energy-efficient.

## 1. Introduction

Spiking recurrent neural networks (SRNNs) are a direct analogue of biological neural networks (Maass, 1997) and can achieve relatively good performance on complex tasks as compared to classic RNN architectures (gated recurrent units, long short-term memory) or feedforward artificial neural networks (ANNs) (Bellec et al., 2018; Yin et al., 2021; Fang et al., 2021). SRNNs also offer energy-efficient solutions for neuromorphic computing (Roy et al., 2019). The performance of SRNNs has been improved due to several developments, including: the use of surrogate gradients to implement back-propagation-through-time (BPTT) (Werbos, 2002); the use of spiking neurons with adaptation mechanisms (Bellec et al., 2018); and increasing or optimizing the heterogeneity across neurons (Perez-Nieves et al., 2021; Bittar and Garner, 2022). However, many properties of biological neurons remain to be implemented in SRNNs, and the way in which these properties affect the performance and energy efficiency of SRNNs remains partially unexplored.

Several studies indicate that the performance of SRNNs can be significantly boosted by using neurons with adaptive thresholds or currents, as compared to the standard LIF models (Bellec et al., 2020, Bellec et al., 2018; Baronig et al., 2025; Bittar and Garner, 2022). Networks with adapLIFs (adaptive Linear Integrate Fire neurons) are able to reproduce both sub-threshold behaviour of so-called excitability class 1 and class 2 neurons (Baronig et al., 2025; Hodgkin, 1948; Izhikevich, 2000), while LIF neurons are restricted to class 1 neurons. Furthermore, these neurons show resonance functions giving rise to subthreshold oscillatory dynamics, and they add longer memory time constants allowing to solve tasks with more memory requirements. The addition of such neurons can be seen as an example of boosting performance by introducing heterogeneity (Perez-Nieves et al., 2021).

However, the implementation of adapLIF neurons in a task-solving SRNN (Bittar and Garner, 2022; Baronig et al., 2025) still leads to voltage dynamics with major fluctuations that are not biologically plausible. Recent work suggests that clipping a neuron’s voltage distribution can improve the performance of a feedforward SNN (Guo et al., 2022). In biological neurons, voltage dynamics are constrained by specific mechanisms, namely the existence of reversal potentials. Reversal potentials lead to voltage-dependent synaptic currents, and can in fact reverse the sign of GABAergic and glutamatergic synaptic inputs. For example, GABAergic inputs become excitatory when the receiving neuron’s voltage is below the chloride reversal potential. Here, we define a new neuron model in which GABAergic and glutamatergic neurons give rise to recurrent synaptic inputs that are governed by a reversal potential. The SRNN thus contains two classes of neurons with positive and negative reversal potentials, mimicking Dale’s law (i.e. neurons using a single neurotransmitter), which are inhibitory or excitatory depending on the receiver neuron’s membrane potential.

We performed simulations on the benchmark Spiking Heidelberg Digits (SHD) (Cramer et al., 2022) classification task and evaluated our biologically inspired model comparing it to the state-of-the-art neuron model adapLIF. We provide a theoretical analyses of the model’s dynamics and analyses of the subthreshold resonance of the adapLIF neurons. We show that introducing a reversal potential boosts task performance and energy efficiency of the SRNN.

## 2. Results

Our starting point is the well-known adapLIF neuron model (Baronig et al., 2025; Bittar and Garner, 2022; Bellec et al., 2018) which we defined in our *baseline model*. We used used a symplectic-Euler (SE) discretization adapted from (Baronig et al., 2025) described in the Methods section,

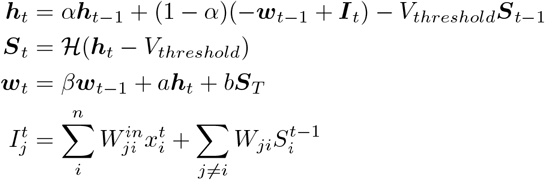

Note that using the bounds on the parameters *α, β, a* and *b* as in (Bittar and Garner, 2022), the trained SRNN contained neurons whose membrane potentials were hypersensitive to the first external input value and diverge to either positive, or negative, infinities. We prevented this effect by using the bound *a ≥* 0 as in (Baronig et al., 2025); note that this bound is also required for neurons to develop sub-threshold resonance (see Methods). We compared the *baseline model* with two other models in which aspects of biological networks / neurons were introduced:

- *Dale model*, which uses the same equations above but with a restriction on the recurrence matrices to obey Dale’s law;
- *AM+GA model*, which uses Dale’s law and in addition implements the voltage reversal potential, such that synaptic inputs are modulated based on the post-synaptic neuron’s membrane potential (Figure1).

**Figure 1.**
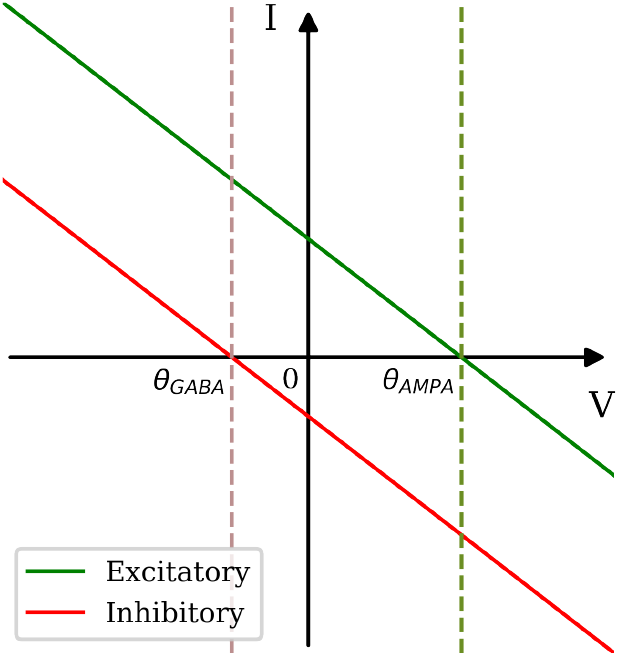
Input gains for excitatory (Green) and inhibitory (Red) pre-synaptic signals based on post-synaptic neuron’s membrane potential. *θ* represents the threshold for the respective glutamergic/GABAergic voltage reversal potential

The voltage reversal potential was mathematically described by changing the input current, ***I***, in the *baseline model* to:

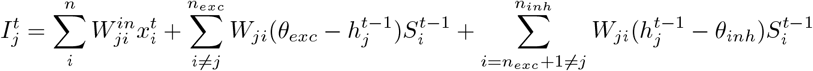

Here, *θ*_*exc*_ and *θ*_*inh*_ are the reversal potentials analogous to glutamatergic (AMPA) and GABAergic inputs, respectively. To determine the relative contributions of these “glutamateric” and “GABAergic” reversal potentials, we further defined two additional models: *AMPA, GABA* where we implemented the voltage reversal potential either for glutamergic or GABAergic inputs (i.e. only the green/red line in Figure1).

Next, we tested the models on the standard Spiking Heidelberg Digits (SHD) classification task. The spiking RNNs were trained offline with supervised learning for two different-sized networks: Architecture 1, which had one hidden layer (and a readout layer); and architecture 2, which contained two hidden layers ( and a readout layer). The SRNNs were trained via BPTT (see Methods). We evaluated task performance on the test set, through <u>accuracy</u>; and energy consumption, through two measures, which describe practically useful energy measures when implementing these networks: <u>Scaled Median Absolute Deviation (Scaled MAD)</u> and <u>Spike energy consumption</u>. These energy measures estimate the average energy consumption per trial by weighing the mean network activity, 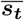 and the mean synaptic transmission of the network caused by spiking activity activity,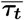. In more detail, the latter measure can be described for a particular sample m and time point t:

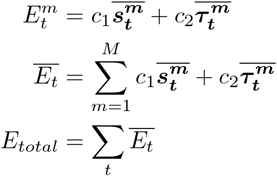

where

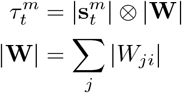

Here, *c*_1_, *c*_2_ are constants that weigh the impact of activity and synaptic transmission (following Ali et al. (2022)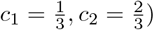. We can inversely correlate this measure with network sparseness and thus with the network’s energy efficiency (see Methods for details). Table 1 summarizes the accuracy and energy efficiency of the different models.

**Table 1:** Results for different models for two different size networks. The number of layers on the left refers to the number of hidden layers in the trained networks. *Accuracy* and *Voltage fluctuation*, relate to performance on the test set averaged across 5 iterations of the model; *Voltage fluctuation* measure is the Scaled MAD.

|  | Model | Accuracy(%) | Test Spike Energy | Voltage fluctuations |
| --- | --- | --- | --- | --- |
| 1 Layer | Baseline, a0 | $88.93 \pm 0.62$ | $516.38 \pm 6.79$ | $0.7756 \pm 0.1625$ |
| | Dale | $88.76 \pm 1.40$ | $450.44 \pm 22.35$ | $0.7538 \pm 0.0394$ |
| | **reversal AM+GA** | $88.56 \pm 0.91$ | **$131.33 \pm 3.27$** | **$0.4198 \pm 0.0511$** |
| | reversal GABA | $89.03 \pm 0.96$ | $292.33 \pm 12.16$ | $0.5301 \pm 0.0389$ |
| | reversal AMPA | **$89.83 \pm 0.80$** | $255.23 \pm 9.27$ | $0.6699 \pm 0.0215$ |
| 2 Layers | Baseline a0 | $85.91 \pm 0.83$ | $407.65 \pm 14.10$ | $1.1015 \pm 0.0145$ |
| | Dale | $84.13 \pm 2.48$ | $170.38 \pm 12.52$ | $2.5906 \pm 1.2023$ |
|  | **reversal AM+GA** | **$90.43 \pm 0.96$** | **$121.18 \pm 11.48$** | **$0.8503 \pm 0.0603$** |
| | reversal GABA | $86.35 \pm 1.23$ | $202.17 \pm 11.92$ | $0.8868 \pm 0.0731$ |
| | reversal AMPA | $89.34 \pm 1.25$ | $260.72 \pm 9.50$ | $0.9901 \pm 0.0635$ |

In comparison to the baseline model, the *Dale model* showed a reduction in spike energy for both the 1-layer and 2-layer model. However, the voltage fluctuations increased and decreased for the 2-layer and 1-layer model, respectively.

Introducing the voltage reversal potential led to a substantial and systematic reduction in both energy consumption and voltage ranges. In architecture 1, we observed an approximately 4-fold reduction in spiking energy, and a 2-fold reduction in the *Scaled MAD* of the membrane potential fluctuations for the *AM+GA model* as compared to the baseline model. Furthermore, both spiking energy and voltage fluctuations reduced relative to Dale’s model. In architecture 2, we observed similar improvements in energy efficiency. Thus, neurons fired more sparsely and had a narrower distribution of voltages in the model with reversal potentials. Analysis of the separate effects of AMPA and GABA suggests that both tend to reduce spiking energy and voltage fluctuations, but act in an additive manner when combined (Table 1). To illustrate these effects, we show some of the neuron responses (Figure 2) for the *baseline network* with large membrane potential fluctuations, compared to sparser firing and constrained membrane potentials with the voltage reversal potential for the *AM+GA model*.

**Figure 2.**
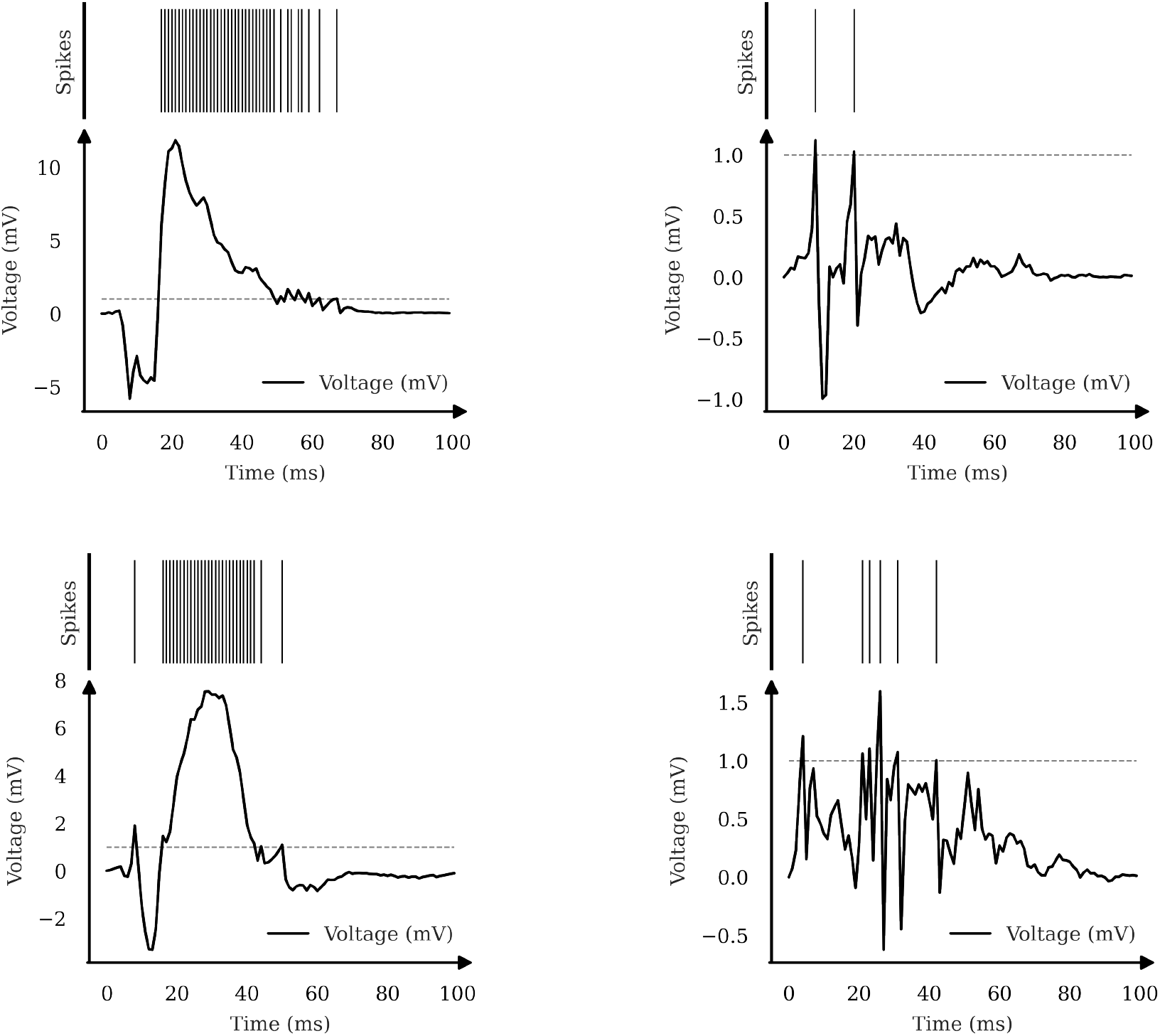
On the left panels we have examples of non biologically constrained model’s neuron profiles for 1 trial. On the right panels we have the same but this time for the model constrained with Dale’s law and reversal potentials

Next, we examined the effect on task performance. Introducing the *Dale model* caused a small, non-significant decrease in accuracy as compared to the baseline model. Strikingly, however, we found a substantial increase in performance from 85% *→* 90% (*p* < 0.001, Two-tailed Welch’s t-test), for the *AM+GA model* as compared to the baseline model and Dale’s law model for the 2-layer network. Analysis of the performances suggest that this performance improvement is primarily due to the inclusion of the AMPA reversal potential, rather than the GABA reversal potential. That is, the performance of the network with only the AMPA reversal potential was comparable to the performance with both the AMPA and GABA reversal potential.

### Analysis of unit properties

To understand how the reversal potentials changes the activity of the neurons in the network, we performed further theoretical analyses of the adapLIF neurons with reversal potentials. As previously shown (Baronig et al., 2025), adapLIF neurons can exhibit resonances at different frequencies, which simply results from the fact that it comprises a system of two coupled differential equations, which therefore can exhibit subthreshold resonance. The performance gain observed with adapLIF neurons likely relates to increased heterogeneity and time constants (Bellec et al., 2018). However, subthreshold resonance may also play a role because RNNs with units configured as damped harmonic oscillators, (CoRNNs - coupled oscillator RNN) have potential theoretical benefits and enhance performance on various tasks (Rusch and Mishra, 2021), in particular when units show resonance at the frequency of the external input (Effenberger et al., 2025).

By analyzing neural properties (see Methods), we found that neurons showed a broad distribution of oscillatory frequencies from about 0 to 8Hz, suggesting the network learns substantially heterogeneous resonance responses. The AM-GA model with reversal potentials had substantially fewer oscillatory units than the Dale’s model and also fewer oscillatory units than the baseline model, with lower eigenvalue magnitudes (i.e. weaker oscillatory responses), and lower frequencies (see Table 2). Notably the AM-GA model also exhibited less adaptation in response to spiking, with smaller values of *b*.

**Table 2:** Parameters across neurons for the different models after training, with standard errors of the mean into:

|  | Parameters | AM+GA | Dale | Baseline |
| --- | --- | --- | --- | --- |
| 1 Layer | $\alpha$ | $0.842 \pm 0.005$ | $0.877 \pm 0.005$ | $0.848 \pm 0.003$ |
| | $\beta$ | $0.985 \pm 0.001$ | $0.983 \pm 0.002$ | $0.985 \pm 0.001$ |
| | $a$ | $0.270 \pm 0.033$ | $0.430 \pm 0.015$ | $0.296 \pm 0.010$ |
| | $b$ | $0.504 \pm 0.041$ | $0.871 \pm 0.069$ | $0.580 \pm 0.019$ |
| | $\vert \lambda\vert $ | $0.911 \pm 0.003$ | $0.928 \pm 0.003$ | $0.913 \pm 0.002$ |
| | Frequency [Hz] | $2.447 \pm 0.2345$ | $3.225 \pm 0.05090$ | $2.709 \pm 0.02879$ |
| | Fraction Oscillatory | $0.703 \pm 0.055$ | $0.931 \pm 0.032$ | $0.808 \pm 0.034$ |
| 2 Layers | $\alpha$ | $0.846 \pm 0.004$ | $0.869 \pm 0.012$ | $0.841 \pm 0.002$ |
| | $\beta$ | $0.985 \pm 0.001$ | $0.987 \pm 0.001$ | $0.985 \pm 0.001$ |
| | $a$ | $0.382 \pm 0.011$ | $0.444 \pm 0.029$ | $0.372 \pm 0.017$ |
| | $b$ | $0.317 \pm 0.048$ | $0.689 \pm 0.044$ | $0.372 \pm 0.017$ |
| | $\vert \lambda\vert $ | $0.913 \pm 0.002$ | $0.926 \pm 0.006$ | $0.910 \pm 0.002$ |
| | Frequency [Hz] | $3.159 \pm 0.1184$ | $3.434 \pm 0.2845$ | $3.647 \pm 0.1253$ |
| | Fraction Oscillatory | $0.789 \pm 0.033$ | $0.922 \pm 0.035$ | $0.894 \pm 0.013$ |

It can be shown that the inclusion of a reversal potential makes the properties of the neuron itself dynamic and dependent on the time-varying input, rather than static. To see this, we note that our system of equations is written as

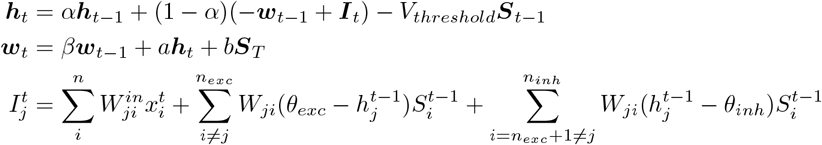

If we consider only one neuron *j* and it is under subthreshold dynamics we can simplify these equations into:

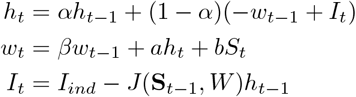

Here, 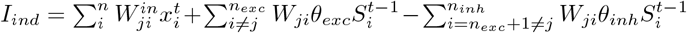 has the feedforward component and the recurrent component independent of *h*_*j*_ and 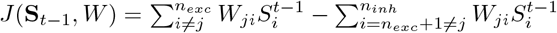 represents the recurrent gain dependent on the membrane potential. Note that *J* > 0 and that *J* represents the sum of the magnitudes of excitatory and inhibitory conductances. We can then write the state transition matrix by plugging in *h*_*t*_ *→ w*_*t*_:

Here, we can see that the inputs themselves determine the state transition matrix *A*, making the response property of the neuron input-dependent, where

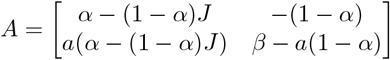

To understand the impact of *J* (amount of conductance) we perform eigenvalue analysis in the subtreshold regime (see Methods for exact derivations). This analysis shows oscillation magnitudes and frequencies that are dependent on recurrent network activity itself, as it determines *J* as a function of time for each neuron.

The expression for the eigenvalue magnitudes equals (see Methods)

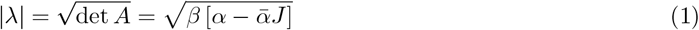

Thus, increased conductances *J* strictly decrease the magnitude of subthreshold oscillations, because *β* > 0 and 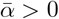.This reduction in amplitude is related to the classic shunting effect of synaptic conductances in biophysics (Koch, 2004).

The eigenvalues are complex as long as

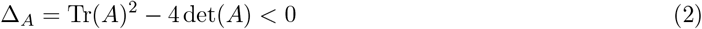

The discriminant is an upward parabola as a function of *J*, reaching a minimum at *J* ^*∗*^, i.e. the system is most likely to oscillate at this point. Oscillatory behavior requires *J* to lie in the interval *{J*_−_, *J*_+_*}*, where

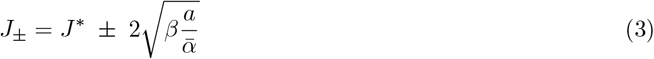

and

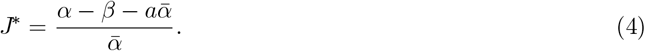

We find in the trained model that all of the neurons in the trained model have *J*^∗^ < 0. Hence, increased conductance *J* invariantly decreases the oscillation magnitude; the eigenvalue magnitudes keep decreasing until the system is not anymore oscillatory. Thus, the best performing model AM+GA already has fewer oscillatory units for *J* = 0, but the tendency to oscillate further decreases with increased *J*. Note, that it is required that *a* ≥ 0 for the system to exhibit oscillations (see Methods). As *a* → 0 the window of conductances *J* in which oscillations occur becomes increasingly narrow.

Finally, for complex discrete eigenvalues we have a frequency 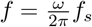 with 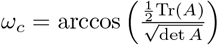(note that deriving the frequency from continuous dynamics as done in other work is inaccurate). We obtain that the maximum frequency is obtained for

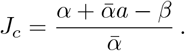

As a consequence, initially frequencies may slightly increase as a function of *J*, until *J*_*c*_, after which they decrease, until the system stops oscillating at *J* ^+^, at which point the frequencies are zero. For our range of parameters, on average *J*_*c*_ ≤ 0 (see Table 2), such that frequencies initially increase (for a subset of units) and then decrease with increased conductance *J*. In sum, the AM+GA model with reversal potentials has fewer oscillatory units with weaker oscillations, and lower frequencies. The reversal potential further leads to oscillations becoming weaker and frequencies showing a non-linear dependence on the conductance *J*.

## 3. Discussion

The performance of SRNNs has been improved due to the introduction of several features, including: the use of surrogate gradients to implement back-propagation-through-time (BPTT) (Werbos, 2002); units showing adaptation mechanisms (Bellec et al., 2018); increased heterogeneity of single-neuron properties (Perez-Nieves et al., 2021; Bittar and Garner, 2022). However, many properties of biological neurons remain to be implemented in SRNNs, and the way in which these properties affect the performance and energy efficiency of SRNNs remain partially unexplored. The implementation of this adapLIF in a task solving RSNN (Bittar and Garner, 2022; Baronig et al., 2025) still leads to non-biologically plausible voltage dynamics as the subthreshold values fluctuate outside of the biological range. In biological neurons, voltage dynamics are constrained by specific mechanisms, namely the existence of reversal potentials (Koch, 2004). Here, we define a new neuron model in which GABAergic and glutamatergic neurons give rise to recurrent synaptic inputs that are governed by a reversal potential. The SRNN thus contains two classes of neurons, mimicking Dale’s law (i.e. neurons using a single neurotransmitter), which are inhibitory or excitatory depending on the receiver neuron’s membrane potential. We trained SRNNs on the Spiking Heidelberg Digits, and show that SRNN with reversal potentials cuts spike energy by up to 4×, while increasing task accuracy. We give a theoretical analysis of the model’s dynamics and analyze the subthreshold resonance of the adapLIF neurons. Increased performance of the model with reversal potential is concomitant with a reduction in the number of oscillatory units and decrease in oscillation frequency as compared to the baseline model. The reversal potential makes subtreshold resonance conductance-dependent, such that increased conductance reduces oscillatory behavior and decreases the frequency.

Further work is needed to precisely identify the reasons why including the reversal potential increases task performance and energy efficiency. There may be several reasons: (1) The reversal potential naturally constrains the voltage range of the neurons, which is explained by the fact that the sign of excitation and inhibition changes around the reversal potential, thereby improving the energy efficiency. This may contribute to task performance by keeping voltages in the range where the surrogate gradients do not vanish, thus mitigating the vanishing gradient problem. In agreement, previous work suggests that clipping a neuron’s voltage distribution can improve the performance of a feedforward SNN (Guo et al., 2022). (2) Mathematically, the reversal potential allows e.g. inhibitory input to have divisive effects on excitatory inputs rather than just subtractive effects, thus enhancing the computational capacity of the network (Koch, 2004). That is, the reversal potential adds a nonlinear integration of inputs inside the neuron which may enhance the computational capacity of neurons. (3) We showed that the reversal potential allows for the resonance frequencies of the individual neurons to become input-dependent. Hence, the resonance function of a neuron may change as a function of the input, which may therefore create more complex and time-dependent feature-selectivities tuned for specific input patterns.

Reversal potentials are, biologically, an unavoidable consequence of the electrochemical gradient of ion channels defined by intracellular and extracellular concentrations. For example, GABA is inhibitory because the intracellular chloride concentration is much lower than the outside of the cell, such that the chemical force is stronger than the electric force when the voltage is above the reversal potential (Koch, 2004). In fact, this is a property that depends on the KCC2 chloride receptor and changes through development, but can also vary between areas (e.g. dentate gyrus) (Ben-Ari, 2007; Sauer et al., 2012). Thus it stands to reason that the precise GABAergic reversal is a property that is evolutionary optimized. In this work, however, we did not optimize the reversal potentials, which may have lead to further increases in task performance and energy efficiency.

In this work we report a performance increase with including Dale’s law with reversal potentials, however there are some other methods that may improve task performance that we did not examine here. Baronig et al. (2025) includes a biologically unrealistic *q* parameter in the adaptation equation (see Methods) that needs to be adjusted for each architecture. They also perform additional preprocessing on the inputs and optimized *α* and *β* for the ADAM Optimizer. We further note that we did not use batch normalization for training here. The reason for this is that batch normalization applied to spiking RNNs does not yield biologically realistic activity.

In sum, the findings here are another step towards more biological realism in spiking neural networks, allowing for efficient and high-performance training with Dale’s law. These findings likely have important applications for neuromorphic computing technologies, in which energy efficiency plays an important role.

## 4. Methods

### Neuron model

We ran a discretized version of the adapLIF model, using the symplectic-Euler (SE) discretization (Baronig et al., 2025). Some small changes to the dynamics were made compared to the (Bittar and Garner, 2022; Baronig et al., 2025) as we will discuss below. The discrete update equations according to the SE-discretization method are:

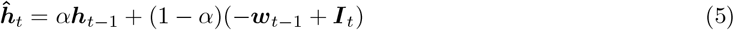

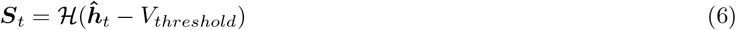

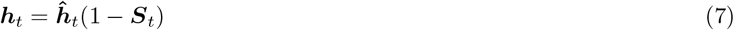

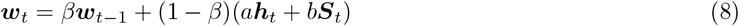

with *h* the hidden membrane potential, *S*_*t*_ a binary variable for the spike output of the neuron, *w* the adaptation variable, and *I* the input currents to the neuron at that time step. The neuron parameters *α, β, a*, and *b* correspond to the membrane decay rate, adaptation decay rate, sub and supra-threshold gain constants, respectively; the parameters remain fixed post learning. We initialize them over a biologically plausible range based on (Bittar and Garner, 2022; Baronig et al., 2025) and allow them to train individually, per neuron, to add heterogeneity to the network (Perez-Nieves et al., 2021).

The discrete update equations we used for the simulations are:

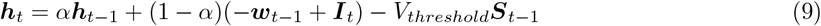

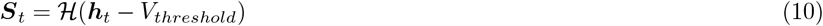

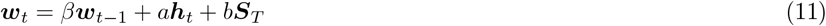

### Differences from previous works

There are two major differences from Eq.(5, 6, 7, 8) to Eq.(9, 10, 11). Firstly, we avoid the hard reset, Eq.(7) used in (Baronig et al., 2025), as this represented in preliminary tests a 7% drop off in accuracy, and use instead a soft reset similar to the one used in (Bittar and Garner, 2022), by adding “ − *V*_*threshold*_***S***_*t*−1_” in Eq.(9).

Secondly, we drop the (1 − *β*) term in Eq.(8) as the network did not learn efficiently, accuracy peaked [55%, 60%] (compared to [85%, 95%] we get otherwise). This is due to (1 − *β*) being very small (in our range of values) effectively removing the dynamical component of our adaptation variable. The scaling of (1 − *β*) was solved in (Bittar and Garner, 2022) identically to us (in the source code, as in the original paper they have that scaling factor present); in (Baronig et al., 2025) by adding another scaling factor *q* reshaping Eq.8 to:

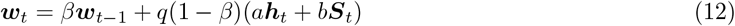

We chose to remove the term from our main results since (Baronig et al., 2025) suggests *q* needs to be tuned for each task as a global gain on the neuron’s adaptation constants.

### Additions to the neuron model

To the baseline equations, Eq.(9, 10, 11), we added two biological realistic features: (1) Dale’s law; and (2) a voltage reversal potential.

1. *Dale’s law* restricts the communication of a neuron to a single neurotransmitter for all postsynaptic connections. This means for our computation that we divide our nodes into two sub-populations with their recurrent weights either positive (excitatory neurons) or negative (inhibitory neurons). The two populations comprise 50% of the population in this case, which was done in order to implement an inductive bias of balanced E/I.
2. By using a *reversal potential*, dendritic inputs to neurons are scaled by its local voltage potential. Its existence is well-established in neuroscientific literature, (Brown et al., 1971; Mofidi et al., 2020; Eisenberg et al., 2015; Koch, 2004).

To model the reversal potential, we changed our input current equation to:

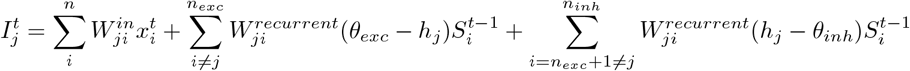

### Theoretical derivations of the neuron model

The system of equations in the subthreshold regime is given by

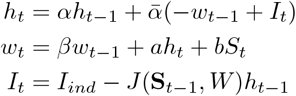

where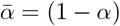. Here, 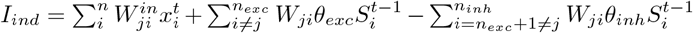, is the feedforward component plus the recurrent component independent of *h*_*j*_.

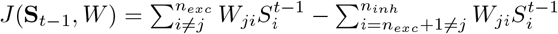 is the recurrent gain dependent on the membrane potential.

Setting *S*_*t*_ = 0 (sub-treshold regime), the system of equations can be written as:

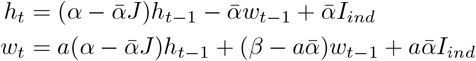

We can then define the subtreshold state transition matrix *A*,

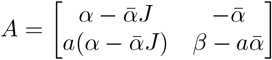

We have

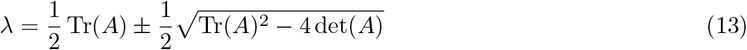

and

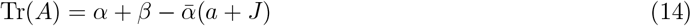

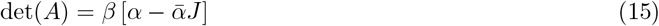

### Eigenvalue magnitude

We first consider the case that there is a pair of complex eigenvalues, i.e. the neuron exhibits subthreshold resonance (which occurs in more than half of the measured neurons). In this case we have the eigenvalue magnitude

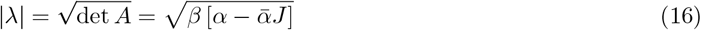

Hence, increased recurrent input conductances *J* tends to decrease the magnitude of subthreshold oscillations as *β* > 0 and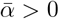.

### Effect of input on tendency to oscillate

The eigenvalues are complex as long as

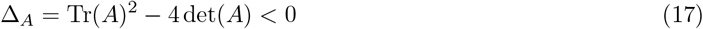

The discriminant is an upward parabola, reaching a minimum at *J* ^*∗*^, i.e. the system is most likely to oscillate at this point, with complex eigenvalues in the interval

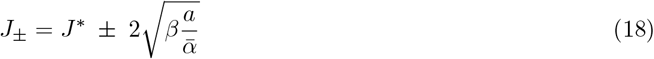

Where

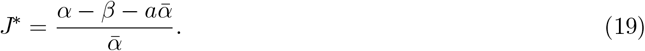

We can see that typically *J* ^*∗*^ < 0 for our range of parameters (Table 2). As the conductance *J* increases, there is a conductance value *J* ^+^ at which point the neuron stops oscillating. Hence, increased *J* leads to decreasing eigenvalue magnitudes, until the neuron falls outside the range of oscillatory behavior.

Increased adaptation *a* widens the window in which oscillations occur, increased membrane potential time constant (*α*) also increases the window. If *a* > 0, there exists an interval *J ∈* (*J*_−_, *J*_+_) in which the eigenvalues are complex, since 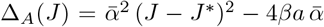 In the absence of an input, *J* = 0, we can see that Δ_*A*_ < 0 if

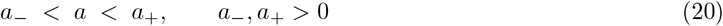

Here

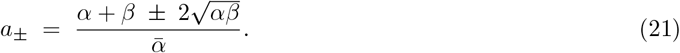

### Bounds on the stability of the system

Stability is given by the Schur inequalities.

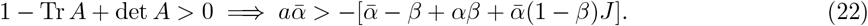

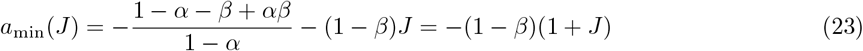

If *a* > 0 and *J* > 0 then *a* > *a*_min_(*J* ). This motivates the bound *a* > 0 to ensure stability.

We have from | det *A*| < 1

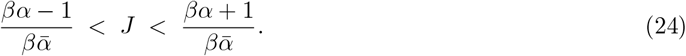

Finally we have

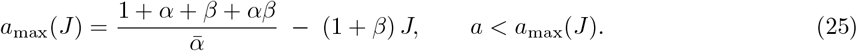

For our ranges of values this implies that *J*_*max*_ is very large (> 10) and thus the system given our parameters is stable. It is further evident that as the network grows larger in size, the network weights should be scaled as 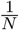 in order to prevent *J* from growing too large.

*J ‘s Critical point*. For complex discrete eigenvalues we have a frequency 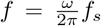 We will derive the dependency of *ω* on *J*. We shall use the correct expression for discrete-time dynamics (note that the continuous time expression would not necessarily be accurate). We define the numerator/denominator of 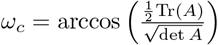 as *g, p*, and the full argument of the arccos as *f*. We are going to be searching for the critical point w.r.t *J*.

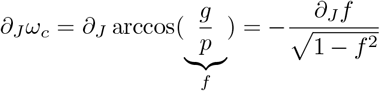

Expanding the terms and simplifying we obtain:

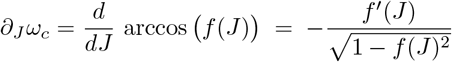

Hence the critical point occurs when *f* ^*′*^(*J* ) = 0. We have

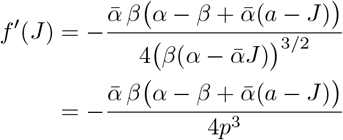

Thus, the critical point occurs when 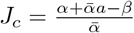.

For the second derivative evaluated at the critical point, we have

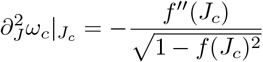

Note that

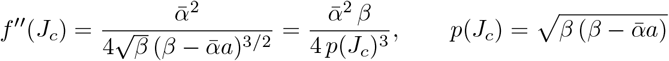

Thus, *f* ^*′′*^(*J*_*c*_) > 0, and the critical point *J*_*c*_ is always a maximum.

Note that we require that *J*_*c*_ ≤ *J* ^+^, i.e. we require that it lies in the domain of complex eigenvalues. Thus, if *J*_*c*_ > 0, initially frequencies increase as a function of *J*, until *J* = *J*_*c*_, after which the frequency decreases; if *J*_*c*_ < 0, frequencies always decrease. At *J* ^+^, we have Tr(*A*)^2^ = 4 det(*A*) and hence *ω*_*c*_ = 0. Hence, frequencies decrease until *ω*_*c*_ = 0 after which sub-threshold oscillations disappear.

### Data

We trained networks on the Spiking Heidelberg Digits (Cramer et al., 2022). This dataset consists of close to 10000 aligned audio recordings of spoken digits, from 0 to 9, in English and German from 12 speakers, where two are exclusive to the test set. These recordings are labelled according to *digit* and *language*, yielding a total of 20 classes. We generate the neural network inputs using the *sparse data generator* function provided by Cramer et al. (2022) which discretizes the spike times based on the number of time steps we provide and an additional argument *max time, T*, which caps the upper limit for the binning time. In our case, we used 100 time steps and *T* = 1. There are 700 input channels, the values of which are integers representing the number of times that a neuron fired in that time bin.

### Task and Neural network architectures

We examined the performance of different neural networks on the ability to correctly learn to classify the SHD data in a supervised setting. We used connected internal feedback spiking recurrent neural network (SRNN) with a final readout layer, similar to (Bittar and Garner, 2022). We used two different architecture, which had the same type of readout layer:

### Architecture 1

Here we used a 2-layer deep network (1 hidden layer + readout layer) with 256 neurons in the hidden layer, with a total number of 250900 trainable parameters. The input currents for each neuron 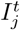 are going to be of the form:

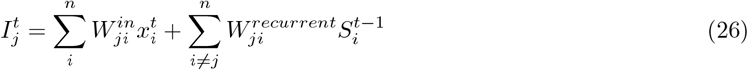

with 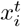 the value for the *i*th input channel. In vector notation we have

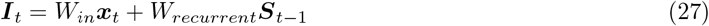

where *W*_*in*_ is the 700×256 input weight matrix and *W*_*recurrent*_ is the 256×256 recurrent weight matrix with diagonal filled with 0s.

### Architecture 2

Here we trained a 3-layer deep network (2 hidden layers + readout layer) with 128 neurons in each of the hidden layers. The total number of trainable parameters was 142356. The vast majority, *≈* 83%, of the difference in the number of trainable parameters comes from the input connections to the network. We can again write the input current for each neuron 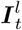 where *l* refers to the hidden layer of the network:

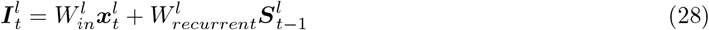

With 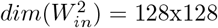 and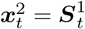.

### Readout layer and loss function

The readout layer of our network consisted of a leaky integrator neuron (LI), achieved by setting *a* = *b* = 0 and removing the reset mechanism. From this output, we would compute the loss function defined in (Bittar and Garner, 2022). For each time step, we apply a softmax function to the hidden potentials of the readout layer across neurons at that time step, then we sum these normalized values across time steps for all neurons and apply a cross-entropy loss with the target class for each trial. For each batch, *n*, the loss was computed as:

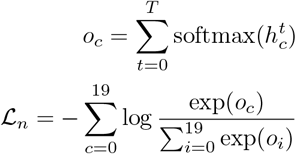

For the reported results we used the same regularization loss function as in (Bittar and Garner, 2022). Additionally, for the 1 hidden layer network we obtained, similar performance using the regularization function from (Zenke and Vogels, 2021) and computing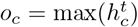.

### Parameter Initialization

We briefly explored the hyperparameter (*α, β, a* and *b*) space and settled on 2 settings, one for each architecture. This space is quite large and since we focused on comparability we decided to base it on previous work with similar models (Bittar and Garner, 2022; Baronig et al., 2025). The hyperparameters were initialized uniformly across the accepted range and during learning were clamped to their bounds to ensure realistic dynamics.

The weights either followed the Xavier initialization, or were sampled from a normal distribution with specified mean and standard deviation using the built in *PyTorch* functions. When using Dale’s law we additionally split the recurrence matrix in half for the excitatory/inhibitory sub-populations. We took the minimum absolute values for the weights (*mavw*) and for the excitatory afferent weights any negative initialized weights were set to *mavw*, and in the inhibitory the symmetric happened.

### Training details

We trained all the models for 100 epochs across the training set of the SHD data. We trained each model for 5 different iterations with the same initialization setup. Every setting used was fixed for each architecture and, for the most part, shared between architectures. These shared settings are specified in table 3. The different settings are specified in Table 4. Two more boolean parameters exist in the dictionaries used, which serve to alter the dynamics of the model: *Apply Dale Law, Use VRP*. For the readout layer the parameters used are specified in Table 5.

**Table 3:** Settings shared for both architectures

| Setting names | Values |
| --- | --- |
| Voltage threshold | 1.0 |
| $\beta$ range | $[e^{-1/3}, e^{-1/120}]$ |
| a range | $[0.0, 1.0]$ |
| b range | $[0.0, 2.0]$ |
| $\theta_{inh}$ | -1.0 |
| $\theta_{exc}$ | 2.0 |
| $\arctan \alpha$ | 2.0 |
| Dropout probability | 0.1 |
| Init $W_{i \rightarrow h}$ with Xavier | True |
| Init $W_{h \rightarrow h}$ with Xavier | True |
| Use adaptable parameters | True |

**Table 4:** Settings specific to each architecture

| Setting names | Architecture 1 | Architecture 2 |
| --- | --- | --- |
| Title | preset1L_params_hidden_Xavier | garnerpaper_params_hidden |
| $\alpha$ range | $[e^{-1/5}, e^{-1/120}]$ | $[e^{-1/5}, e^{-1/25}]$ |

**Table 5:** Readout Layer settings used

| Setting names | Readout layer |
| --- | --- |
| Title | garnerpaper_params_readout |
| Voltage threshold | 1.0 |
| $\alpha$ range | $[e^{-1/5}, e^{-1/25}]$ |
| $\beta$ range | $[e^{-1/3}, e^{-1/120}]$ |
| Use adaptable parameters | True |
| Use spike reset | False |
| Init $W_{h \rightarrow o}$ mean | 0.0 |
| Init $W_{h \rightarrow o}$ std | 0.03 |
| $\arctan \alpha$ | 2.0 |

The setting names are slightly different from the ones used in the dictionary made available in the code for readability purposes. The setting *arctan α* is a scaling variable for the surrogate gradient function used; it was kept the same as in (Bittar and Garner, 2022)

### Analyses

The SHD does not have a predefined validation set so all our results came from using the full test set. For all analyses, we computed these values per iteration of *model X* and then found the mean and standard deviation (sd) for the 5 iterations. Per iteration, we ran through the full test set via our sampling function (in batches) computing the value and saving it into an array; we would perform this loop 10 times and then average the result as our final per iteration value.

For *Accuracy*, we took *y*^*∗*^ = arg max_*c*Σ*t*_ softmax(*h*_*t*_)_*c*_ (c denotes the classification neuron that goes from 0 to 19 and *y*^*∗*^ our guess for the label) and compared it to the label, *y*, of the class for each batch.

For *Test Spike Energy*, we computed the average neuron energy consumption at each time point and summed them for the duration of the stimulus. Our measure was inspired from (Ali et al., 2022), equations 7 and 8. We defined it for a particular sample m and time point t in a batch:

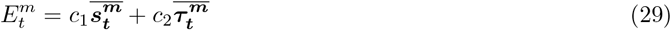

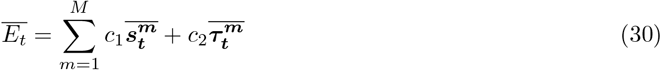

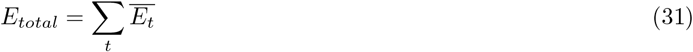

Where

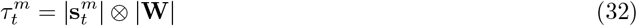

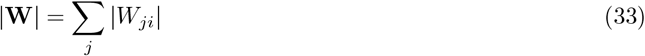

The 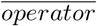 is the average across neurons for a batch until 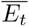. From this point on, it also includes the average across batches. Indeed, one can perform the average across batches or neurons interchangeably, which is how we do it computationally. We chose this notation to maintain with the original notation of (Ali et al., 2022).

This metric has two components for each time step: 1)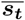 the mean network activity 2) 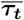 the mean network synaptic transmission caused by said spike activity. Both of these components were averaged across layers to ensure comparability between architectures. This measure is inversely correlated with the sparseness of the network. Since the weight matrices did not differ dramatically between models we can heavily correlate how sparse a network was with how low their energy value was (hence how energy efficient they were).

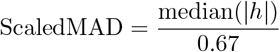

For *Voltage fluctuations*, we used the Scaled median absolute deviation from 0 (Scaled MAD) for the resulting membrane potentials of a batch:

The sampling function has a built-in shuffling property. In combination with the way we loop through the test set, for each iteration value, we are computing a Monte Carlo estimate of the expected measures. However, due to the order of magnitude of the samples our value is statistically valid.

### Bound on a and Reversal potentials

The bounds on the subthreshold adaptation constant, *a*, have a strong impact on the membrane dynamics (Baronig et al., 2025). We bounded *a ≥* 0 according to Baronig et al. (2025) and our own derivations (see Methods). This regime *a ≥* 0 represents the adaptation regime, with increased voltages leading to increased adaptation.

The reversal potential values were based on heuristic estimates, approximately matching reversal potentials in biological neurons, and were not subject to optimization.

## Acknowledgements

This project was financed by DFG VI Grants (908/5-1 and 908/7-1;505660261; 520285844; SPP LOOPS); an NWO VIDI Grant; the Dutch Brain Interface Initiative (DBI2); ERC starting grant (850861) SPATEMP.

## Author Contributions

Funding Acquisition, M.V.; conceptualization, M.P.,,M.V.; simulations, M.P.; mathematical analysis, M.P., C.G., M.V.; writing main draft M.P. and M.V.; supervision, M.V.

## References

Ali, A., Ahmad, N., Groot, E.d., Gerven, M.A.J.v., Kietzmann, T.C., 2022. Predictive coding is a consequence of energy efficiency in recurrent neural networks. Patterns 3. URL: https://www.cell.com/patterns/abstract/S2666-3899(22)00271-9, doi:10.1016/j.patter.2022.100639. publisher: Elsevier.

Baronig, M., Ferrand, R., Sabathiel, S., Legenstein, R., 2025. Advancing Spatio-Temporal Processing in Spiking Neural Networks through Adaptation. URL: http://arxiv.org/abs/2408.07517, doi:10.48550/arXiv.2408.07517. 2408.07517 [cs].

Bellec, G., Salaj, D., Subramoney, A., Legenstein, R., Maass, W., 2018. Long short-term memory and learning-to-learn in networks of spiking neurons. Advances in neural information processing systems 31.

Bellec, G., Scherr, F., Subramoney, A., Hajek, E., Salaj, D., Legenstein, R., Maass, W., 2020. A solution to the learning dilemma for recurrent networks of spiking neurons. Nature Communications 11, 3625, doi:10.1038/s41467-020-17236-y.

Ben-Ari, Y., 2007. Gaba: A pioneer transmitter that excites immature neurons and generates primitive oscillations. Physiological Reviews 87, 1215–1284. doi:10.1152/physrev.00017.2006.

Bittar, A., Garner, P.N., 2022. A surrogate gradient spiking baseline for speech command recognition. Frontiers in Neuroscience 16, 865897. URL: https://www.frontiersin.org/articles/10.3389/fnins.2022.865897/full, doi:10.3389/fnins.2022.865897.

Brown, J.E., Muller, K.J., Murray, G., 1971. Reversal Potential for an Electrophysiological Event Generated by Conductance Changes: Mathematical Analysis. Science 174, 318–318. URL: https://www.science.org/doi/10.1126/science.174.4006.318, doi:10.1126/science.174.4006.318.

Cramer, B., Stradmann, Y., Schemmel, J., Zenke, F., 2022. The heidelberg spiking data sets for the systematic evaluation of spiking neural networks. IEEE Transactions on Neural Networks and Learning Systems 33, 2744–2757. doi:10.1109/TNNLS.2020.3044364.

Effenberger, F., Carvalho, P., Dubinin, I., Singer, W., 2025. The functional role of oscillatory dynamics in neocortical circuits: a computational perspective. Proceedings of the National Academy of Sciences 122, e2412830122.

Eisenberg, B., Liu, W., Xu, H., 2015. Reversal permanent charge and reversal potential: case studies via classical Poisson–Nernst–Planck models. Nonlinearity 28, 103–127. URL: https://iopscience.iop.org/article/10.1088/0951-7715/28/1/103, doi:10.1088/0951-7715/28/1/103.

Fang, W., Yu, Z., Chen, Y., Huang, T., Masquelier, T., Tian, Y., 2021. Deep residual learning in spiking neural networks. Advances in Neural Information Processing Systems 34, 21056–21069.

Guo, Y., Tong, X., Chen, Y., Zhang, L., Liu, X., Ma, Z., Huang, X., 2022. RecDis-SNN: Rectifying Membrane Potential Distribution for Directly Training Spiking Neural Networks, in: 2022 IEEE/CVF Conference on Computer Vision and Pattern Recognition (CVPR), IEEE, New Orleans, LA, USA. pp. 326–335. URL: https://ieeexplore.ieee.org/document/9880053/, doi:10.1109/CVPR52688.2022.00042.

Hodgkin, A.L., 1948. The local electric changes associated with repetitive action in a non-medullated axon. The Journal of physiology 107, 165.

Izhikevich, E.M., 2000. Neural excitability, spiking and bursting. International journal of bifurcation and chaos 10, 1171–1266.

Koch, C., 2004. Biophysics of computation: information processing in single neurons. Oxford university press.

Maass, W., 1997. Networks of spiking neurons: The third generation of neural network models. Neural Networks 10, 1659–1671. URL: https://linkinghub.elsevier.com/retrieve/pii/S0893608097000117, doi:10.1016/S0893-6080(97)00011-7.

Mofidi, H., Eisenberg, B., Liu, W., 2020. Effects of Diffusion Coefficients and Permanent Charge on Reversal Potentials in Ionic Channels. Entropy 22, 325. URL: https://www.mdpi.com/1099-4300/22/3/325, doi:10.3390/e22030325. number: 3 Publisher: Multidisciplinary Digital Publishing Institute.

Perez-Nieves, N., Leung, V.C.H., Dragotti, P.L., Goodman, D.F.M., 2021. Neural heterogeneity promotes robust learning. Nature Communications 12, 5791. URL: https://www.nature.com/articles/s41467-021-26022-3, doi:10.1038/s41467-021-26022-3.

Roy, K., Jaiswal, A., Panda, P., 2019. Towards spike-based machine intelligence with neuromorphic computing. Nature 575, 607–617. URL: https://www.nature.com/articles/s41586-019-1677-2, doi:10.1038/s41586-019-1677-2. publisher: Nature Publishing Group.

Rusch, T.K., Mishra, S., 2021. Coupled oscillatory recurrent neural network (cornn): An accurate and (gradient) stable architecture for learning long time dependencies, in: International Conference on Learning Representations. URL: https://openreview.net/forum?id=F3s69XzWOia.

Sauer, J.F., Strüber, M., Bartos, M., 2012. Distinct subcellular input specificities and plasticity rules for gabaergic synapses of parvalbumin-positive interneurons onto ca1 pyramidal cells and dentate granule cells. Journal of Neuroscience 32, 4224–4239. doi:10.1523/JNEUROSCI.6277-11.2012.

Werbos, P.J., 2002. Backpropagation through time: what it does and how to do it. Proceedings of the IEEE 78, 1550–1560.

Yin, B., Corradi, F., Bohté, S.M., 2021. Accurate and efficient time-domain classification with adaptive spiking recurrent neural networks. Nature Machine Intelligence 3, 905–913. URL: https://www.nature.com/articles/s42256-021-00397-w, doi:10.1038/s42256-021-00397-w. publisher: Nature Publishing Group.

Zenke, F., Vogels, T.P., 2021. The remarkable robustness of surrogate gradient learning for instilling complex function in spiking neural networks. Neural computation 33, 899–925.

